# Role of Solvent Compatibility in the Phase Behavior of Binary Solutions of Weakly Associating Multivalent Polymers

**DOI:** 10.1101/2021.09.09.459431

**Authors:** Jasper J. Michels, Mateusz Brzezinski, Tom Scheidt, Edward A. Lemke, Sapun H. Parekh

**Affiliations:** Max Planck Institute for Polymer Research, Ackermannweg 10, 55128 Mainz, Germany; Institute for Molecular Biology, Johannes Gutenberg University, Ackermannweg 4, 55128 Mainz, Germany; Department of Biomedical Engineering, The University of Texas at Austin, 107 W Dean Keeton Street Stop C0800, Austin TX 78712, USA

## Abstract

Condensate formation of biopolymer solutions, prominently those of various intrinsically disordered proteins (IDPs), is determined by “sticky” interactions between associating residues, multivalently present along the polymer backbone. Using a ternary mean field “stickers-and-spacers” model, we demonstrate that if sticker association is of the order of a few times the thermal energy, a delicate balance between specific binding and non-specific polymer-solvent interactions gives rise to a particularly rich ternary phase behavior under physiological circumstances. For a generic system represented by a solution comprising multi-associative scaffold and client polymers, the difference in solvent compatibility between the polymers modulates the nature of isothermal liquid-liquid phase separation (LLPS) between associative and segregative. The calculations reveal regimes of dualistic phase behavior, where both types of LLPS occur within the same phase diagram, either associated with the presence of multiple miscibility gaps, or a flip in the slope of the tie-lines belonging to a single coexistence region.

## Introduction

A recent paradigm shift has shown important eukaryotic cellular functions to be regulated by biomolecular condensates that form via liquid-liquid phase separation (LLPS) from the cytoplasm or nucleoplasm (1, 2, 3, 4). These “membraneless organelles” (MLOs), of which nucleoli, stress granules, P-bodies and cajal bodies are examples, exist as suspended, dense droplets, typically containing multiple species of biomacromolecules, such as proteins and RNA. In particular, intrinsically disordered proteins (IDPs) are highly prevalent in such condensates, which reflects their pivotal role in driving the LLPS. Besides in MLOs, the intrinsically disordered regions (IDRs) of such proteins may also be part of ordered constructs, such as the nuclear pore complex (NPC). In this case, the IDRs of nucleoporins (Nup) form a crowded assembly that provides for the selective “gateway” properties of the central region of the NPC for nucleocytoplasmatic cargo transport (5, 6, 7, 8), which can be unassisted or assisted by the presence of a nuclear transport receptor (NTR).

Typically, IDPs or IDRs drive LLPS through non-covalent association between specific amino-acid sequences that occur along the backbone in a multivalent fashion (9). The nature of such “sticky interactions” may for instance be hydrogen bonding (10), electrostatic (11), π-π stacking (12), π-cation interaction (13, 14), due to the hydrophobic effect (15, 16, 17) or a combination of binding motifs (10, 18). Although MLOs may easily comprise complex multicomponent mixtures, the phase behavior is often determined by a few or even a single species acting as a “scaffold” to which secondary components associate as “clients” or ligands (19), which, depending on the interaction strength, modulate the driving force for condensate formation (20, 21). Hence, *in vitro* approaches targeting the behavior and LLPS of reduced, experimentally accessible solutions of native (8, 10, 22), mutant (23) or engineered (24, 25, 26) associating biomacromolecules are effective experimental tools in the elucidation of complex cellular mechanisms based on LLPS.

For the same reason, theoretical models for LLPS that consider a limited number (*i.e.* one, two or three) of interacting components (27) are valuable model systems that often provide for predictive context for both *in vivo* and *in vitro* experiments. Specific advantages of such models are their conceptual clarity, computational tractability and the prospect of allowing predictions by analytical theory. Within this context, the “stickers-and-spacers” (SAS) model, as introduced by Semenov and Rubinstein (28, 29, 30), has proven to be a powerful framework in describing and predicting both specific and generic features of the phase behavior of mixtures and solutions of multivalent associating polymers. The SAS concept, which forms the basis for mean-field models (31, 32), as well as coarse-grained simulations (33, 34), parametrizes a multivalent polymer as a chain of sticky residues, separated by segments (“spacers”) comprising non-sticky units. In a typical embodiment of the SAS model, the stickers non-covalently associate with each other to form binary complexes, whereas the spacers contribute to translational entropy. In more complex realizations spacers may also impart correlations between binding events by, for instance, imposing chain rigidity (32, 35).

Of particular theoretical and experimental interest are condensates that comprise two (major) different multivalent biopolymers (24, 36, 37, 38, 39, 40, 41). Recent efforts have focused on cases for which the phase behavior results from heterotypic sticker association, *i.e.* between stickers situated on different polymer species. In the limit of strong binding, *i.e.* for binding energies in excess of about ten times the thermal energy (*kT*), SAS-based models have shown that such associative mixtures to undergo “magic number” (35) and “magic ratio” (42) transitions, based on whether or not the sticker valencies or stoichiometries are commensurate. In many cases, however, sticker binding is significantly weaker (35) and homotypic binding may compete with heterotypic binding. Importantly, if sticker association is relatively weak, *e.g.* of the order of a few times *kT*, it can no longer be regarded as the sole or dominant mechanism underlying the phase behavior of the solution. It is intuitive that in this case the polymer-solvent interactions may significantly contribute to shaping the phase diagram. Unfortunately, studies focusing on the interplay between sticker association and solvent interaction in determining the ternary phase behavior are sparse.

In this work, we use a ternary mean-field SAS-based model to reveal that rich phase behavior arises if contributions from isotropic polymer-solvent interactions and sticker association are of comparable overall magnitude. We calculate isothermal ternary phase diagrams (binodal compositions) for mixtures of two multivalent polymers A and B in a solvent, considering heterotypic (AB) and homotypic (AA and BB) sticker association, as well as non-specific and non-saturating mutual interactions between the two polymers and the solvent. We represent the latter by binary Flory interaction parameters (*χ*_*ij*_) (43). As such, the model includes six types of interactions: three specific and three non-specific, the latter including a non-specific exchange interaction between the polymers. We parametrize the non-specific interactions such that LLPS would not occur if sticker binding were absent.

After describing the model, we define the central case of this work, where Polymer-A acts as a scaffold that, on account of the AA sticker binding, phase separates in its homopolymer solution. In contrast, a homopolymer solution of Polymer-B does not exhibit LLPS, as homotypic B-sticker association is significantly weaker than AA binding. Nevertheless, the “client” Polymer-B acts as a regulator and modulates the propensity of the ternary mixture Polymer-A:Polymer-B:solvent to phase separate. However, we demonstrate that these regulating properties are not only determined by the relative strength of homotypic versus heterotypic sticker binding, but strongly influenced by the difference in solvent compatibility of the polymers. The interplay between the plurality of interactions modulates the nature of the LLPS between *associative*, where the two polymers tend to collect in a concentrated phase and *segregative*, for which the coexisting phases are enriched in either polymer (44, 45). We locate the occurrence of these regimes in the parameter space and highlight cases exhibiting complex, dualistic behavior, such as “double reentrance” and the occurrence of both associative and segregative LLPS within the same phase diagram or even the same miscibility gap.

## Results and discussion

### A stickers-and-spacers model for a binary solution of associating polymers

We formulate a ternary mean-field SAS model that besides non-covalent association between sticky residues on two different polymer species, includes weak, “non-specific” interactions between any neighboring (but unconnected) monomeric units or solvent molecules (see Figure 1). As in the original monopolymer model by Semenov and Rubinstein (28), sticker association is “specific” in that it occurs between designated sites on the polymer chains, but not “orientational” in the sense that structural rigidity limits the directional freedom of the non-covalent bonds, as for instance seen for patchy globular species (46) and folded amino acid sequences (47). Although the solution medium for biopolymers is typically water or an aqueous buffer, we utilize the term “solvent” for the sake of generality.

**Figure 1.**
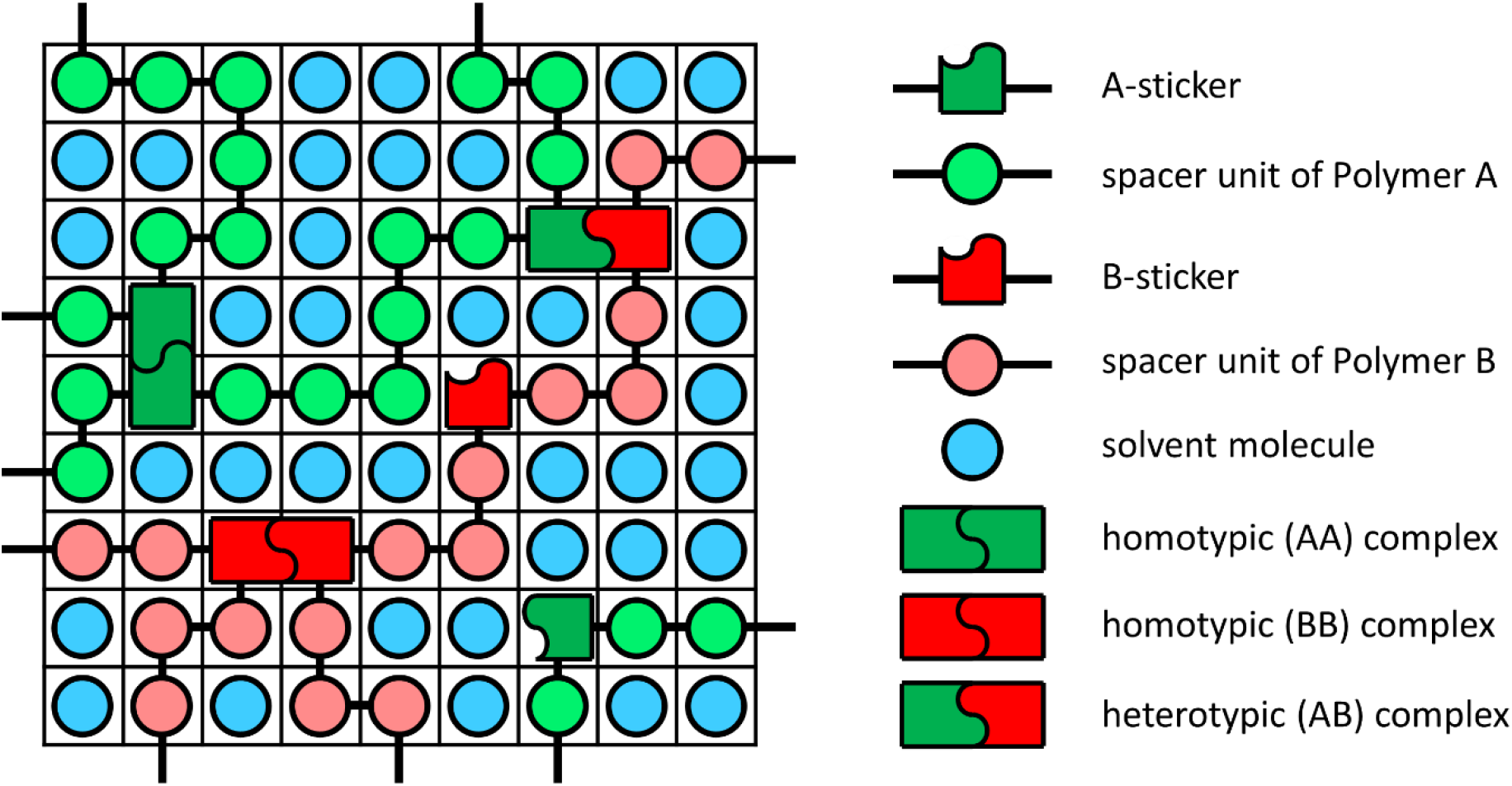
Lattice representation of the ternary mean field stickers-and-spacers model used in this work. Besides the three possible sticker complexes depicted on the right, the monomeric units of the two polymers and the solvent interact through “non-specific” nearest neighbor interactions expressed by binary Flory interaction parameters. For this, no discrimination is made between sticker and spacer monomers.

To predict how the interplay between sticker association, non-specific interaction and chain length determines the phase diagram, we extend the classical Flory-Huggins (*FH*) mixing free energy for a ternary mixture of Polymer-A, Polymer-B and solvent, as we used in previous studies (48, 49), by a contribution from the presence and association of/between stickers (*ST*):

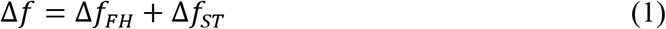

The dimensionless Flory-Huggins free energy density is given by (43):

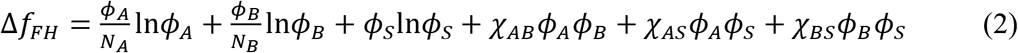

with *ϕ*_*i*_ the volume fraction, *N*_*i*_ the effective molecular size relative to that of a solvent molecule and *χ*_*ij*_ the binary Flory parameters that represent the non-specific polymer-polymer and polymer-solvent exchange interactions. The subscripts A, B and S respectively refer to the two multivalent associative polymers and the solvent. The model is subject to the assumption of incompressibility: *ϕ*_*A*_ + *ϕ*_*B*_ + *ϕ*_*S*_ = 1.

The free energy contribution due to sticker binding is an extension of the original single-species associating polymer model (28), where we assume stickers capable of forming homotypic, *i.e.* AA and BB, as well as heterotypic (AB) non-covalent complexes (see Figure 1). Under the condition that sticker binding is at equilibrium (28), the contribution to the mixing free energy is given by:

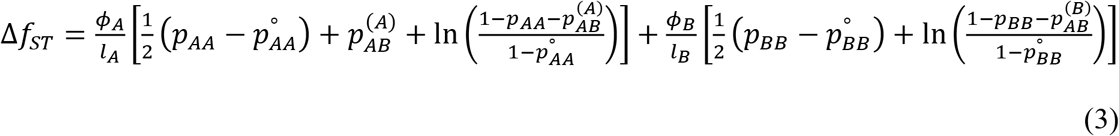

Here, *l*_*i*_ represents the average length of the spacer units in terms of the number of effective monomers and *p*_*ij*_ the fraction of stickers accommodated in the complexes indicated by the corresponding subscript. Specifically, 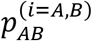 represents the fraction of *i*-type stickers involved in a heterotypic (AB) association. The heterotypic fractions are related according to: 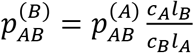, with *C*_*i*=*A,B*_ the number density of *i*-monomers. The sticker contribution is equivalent to the binary model proposed by Olvera de la Cruz *et al.* (31), though here placed in the context of a ternary mixture with *ϕ*_*B*_ becoming an additional independent volume fraction. Furthermore, to be consistent with the Flory-Huggins contribution, the sticker free energy has been formulated relative to that of the pure components, where 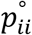 represents the equilibrium fraction of bound stickers in the pure state. Since the terms containing 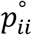 are zeroth or first order in the composition, they do not affect the phase diagram. For completeness, we have given the derivation of the model in Section S1 of the Supplementary Information (SI).

The assumption that sticker binding is in equilibrium yields the relation between the fractions of bound stickers, the binding energies *ε*_*ij*_ and molar association constants *K*_*ij*_:

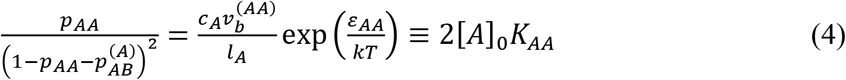

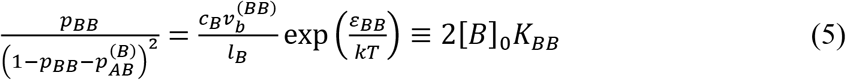

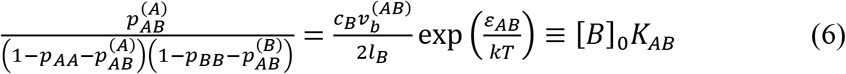

Here, 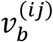 is the volume of a non-covalent bond. [*A*]_0_ and [*B*]_0_ are the total molar concentrations of A and B stickers, *e.g.* expressed in mol l^−1^. The temperature dependence of the sticker association strength is included by defining the enthalpy (Δ*h*_*ij*_) and entropy (Δ*s*_*ij*_) of formation of an *ij*-complex, with: *ε*_*ij*_ (*T*) = Δ*h*_*ij*_ − *T*Δ*s*_*ij*_. As the investigated temperature range is physiologically relevant and hence small (see below), Δ*h*_*ij*_ and Δ*s*_*ij*_ are considered constant. The Floy interaction parameters, which quantify the non-specific interactions, simply scale as ~*T*^−1^ (43), where we treat 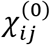 at *T*_ref_ = 273 K as input and reference, so: 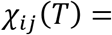 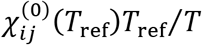.

### Case definition and model parametrization

To demonstrate the consequences and importance of solvent interactions in modulating the phase behavior of a solution of two multivalent associating (bio)polymers, we study a general case based on a scaffold or “driver” (50) polymer (Polymer-A) of which the pure solution phase separates and a second polymer (Polymer-B) that does not phase separate in solution. We refer to Polymer-B as a “regulator” (50) or client, rather than a second scaffold species (51). Another important choice is for homotypic (AA) sticker association, rather than non-specific polymer-solvent interaction, to be the primary driving force for LLPS of a solution of the scaffold Polymer-A. This complies with experimental observations and at the same time provides for the opportunity to show that the ternary phase behavior can be significantly affected even by small variations in generally favorable polymer-solvent interactions.

Table 1 lists the model input parameters that define the above-described case. We assume the scaffold polymer to be larger than the regulator (*N*_*A*_ > *N*_*B*_) but with effective degrees of polymerization within the same order of magnitude. The values for *l*_*i*=*A,B*_ correspond to 50 A-sticker sites on a chain of Polymer-A and ~17 B-stickers on a chain of Polymer-B. To reduce complexity and place emphasis on the effect of polymer-solvent interaction, we set *χ*_*AB*_ to zero in all calculations. The values of 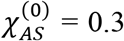 and 0.55 for the non-specific interaction between Polymer-A and the solvent (*χ*_*AS*_) (roughly) represent “marginal” (52) and *θ*-conditions, respectively. The solvent interaction parameter for Polymer-B (*χ*_*BS*_) is scanned in the given range, which corresponds to: 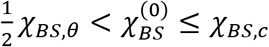, with *χ*_*BS*,*θ*_ and *χ*_*BS*,*c*_ the theta- and critical value (0.5 and ~0.57), the latter defined as: 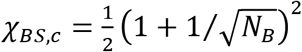. We purposely evaluate *χ*_*AS*_ and *χ*_*BS*_ sufficiently low to exclude LLPS of the homopolymer solutions in the absence of homotypic sticker association. However, since due to their inhomogeneous primary structure many IDPs are not necessarily ideally accommodated in an aqueous environment, we avoid the athermal limit (*χ*_*iS*_ = 0) by a significant margin and hence omit effects related with strong excluded volume contributions.

**Table 1.** Input parameters as used in our study

|  |  |  |  |
| --- | --- | --- | --- |
| $N_A$ | 500 | $\Delta H_{AA}$ (kJ/mol)<sup>a)</sup> | 22.86 |
| $N_B$ | 250 | $\Delta S_{AA}$ (J/molK)<sup>b)</sup> | 113.1 |
| $N_S$ | 1 | $\Delta H_{BB}$ (kJ/mol) | 0 |
| $l_A$ | 10 | $\Delta S_{BB}$ (J/molK) | 0.8314 |
| $l_B$ | 15 | $\Delta H_{AB}$ (kJ/mol) | 5.737 |
| $\chi_{AB}^{(0)}$ | 0 | $\Delta S_{AB}$ (J/molK) | 52.38 – 64.02 |
| $\chi_{AS}^{(0)}$ | 0.3; 0.55 | $T$ (K) | 300 |
| $\chi_{BS}^{(0)}$ | 0.25 – 0.58 | | |

<sup>a)</sup> $\Delta H_{ij} = \mathcal{N}_{Av} \Delta h_{ij}$ ; <sup>b)</sup> $\Delta S_{ij} = \mathcal{N}_{Av} \Delta s_{ij}$ ; $\mathcal{N}_{Av}$ = Avogadro's number

The two rightmost columns of Table 1 parametrize the strength of homotypic and heterotypic sticker association in terms of the binding enthalpies and entropies. The positive values for the enthalpy and entropy of homotypic A-sticker association imply that the LLPS of the homopolymer solution of the scaffold is entropy-driven and therefore exhibits a lower critical solution temperature (LCST), as we shall see below. Such phase behavior is for instance expected if sticker binding occurs via hydrophobic interactions as seen for various elastin-like polypeptides (53, 54, 55, 56), some resilin-like polypeptides, such as An16 resilin (57), as well as the ubiquitin-binding shuttle protein UBQLN2 (18, 58). We also would like to draw a comparison with the phase behavior of FG-Nucleoporins (Nup) (59, 60, 61). We note however, that the choice for sticker binding being entropy- or enthalpy-driven does on a general level not affect the results and conclusions of the (near-)isothermal calculations of the ternary binodals.

To be physically consistent with the driving force for LLPS of the scaffold, homotypic B-sticker association and heterotypic (AB) sticker binding are also entropy driven, although we assume the former very weak. To probe the contribution of sticker association, we scan the strength of heterotypic (AB) binding relative to that of homotypic binding. Doing this by modulating Δ*s*_*AB*_ (see range in Table 1) is in the first place instrumental, – we could also have adjusted the binding strength by varying Δ*H*_*AB*_, – but can be physically interpreted as expressing a variation in the number of solvating water molecules being expelled to the bulk solution upon association between hydrophobic sticker units. The strength of the AA complexes exhibits a steeper temperature dependence than the binding strength of the AB complexes because of the higher enthalpic penalty (see Table 1). Both complexes become stronger with increasing temperature, which forms the basis for the LCST behavior of the scaffold solution. Homotypic B-sticker association is temperature invariant as we set Δ*H*_*BB*_ = 0. The temperature dependence of the binding strength of all possible sticker complexes is expressed by Figure 2, where we have transformed the binding enthalpies and entropies into energies, scaled by *kT*. Following Olvera de la Cruz *et al.* (31), we quantify the relative strength of heterotypic binding by introducing a dimensionless exchange energy, similar to a Flory parameter: 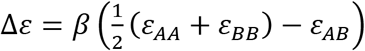, with *β* = 1/*kT*. In view of the fact that the sign of *ε*_*ij*_ is negative, indicating an attractive force, Δ*ε* is positive and increases with increasing AB association strength (see Figure 2).

**Figure 2.**
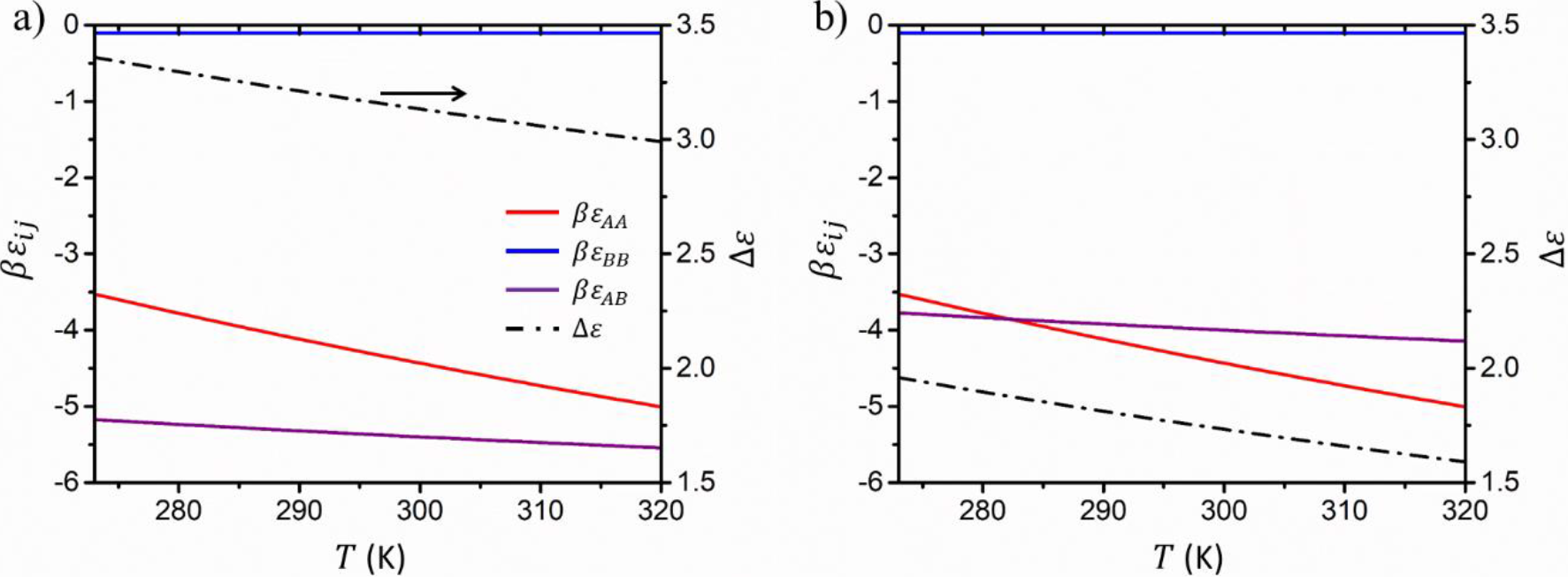
Sticker binding energy as a function of the absolute temperature for strong (a) and weak (b) heterotypic (AB) stickler binding. The purple curves in a) and b) respectively correspond to the upper and lower extreme for Δ*s*_*AB*_ as given in Table 1.

The starting point of our calculations is the binary temperature-composition diagram for a solution of the pure scaffold Polymer-A (Figure 3). The binodal, stability limit (spinodal) and critical point have respectively been numerically obtained in the usual way via the constraints of, respectively, i) equal osmotic pressure and exchange chemical potential in coexisting phases, ii) zero free energy curvature 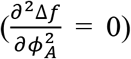 and iii) 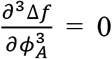. Figure 3a demonstrates the LCST behavior for 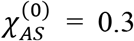. At a sufficiently low, but clearly non-physiological temperature of *T* << 273 K, we expect a second coexistence region exhibiting upper critical solution temperature (UCST), as *χ*_*AS*_ will eventually exceed its critical value. For 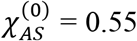 the phase diagram changes drastically and adopts an hour-glass shape and lacks a critical point (Figure 3b) because the two coexistence regions merge. Due to the large difference in molecular size between Polymer-A and the solvent, the phase diagrams are highly asymmetric. Under experimental or physiological conditions the concentrated branch of the binodal curve typically represents the scaffold-rich droplet phase, whereas the dilute branch gives the composition of the surrounding solution.

**Figure 3.**
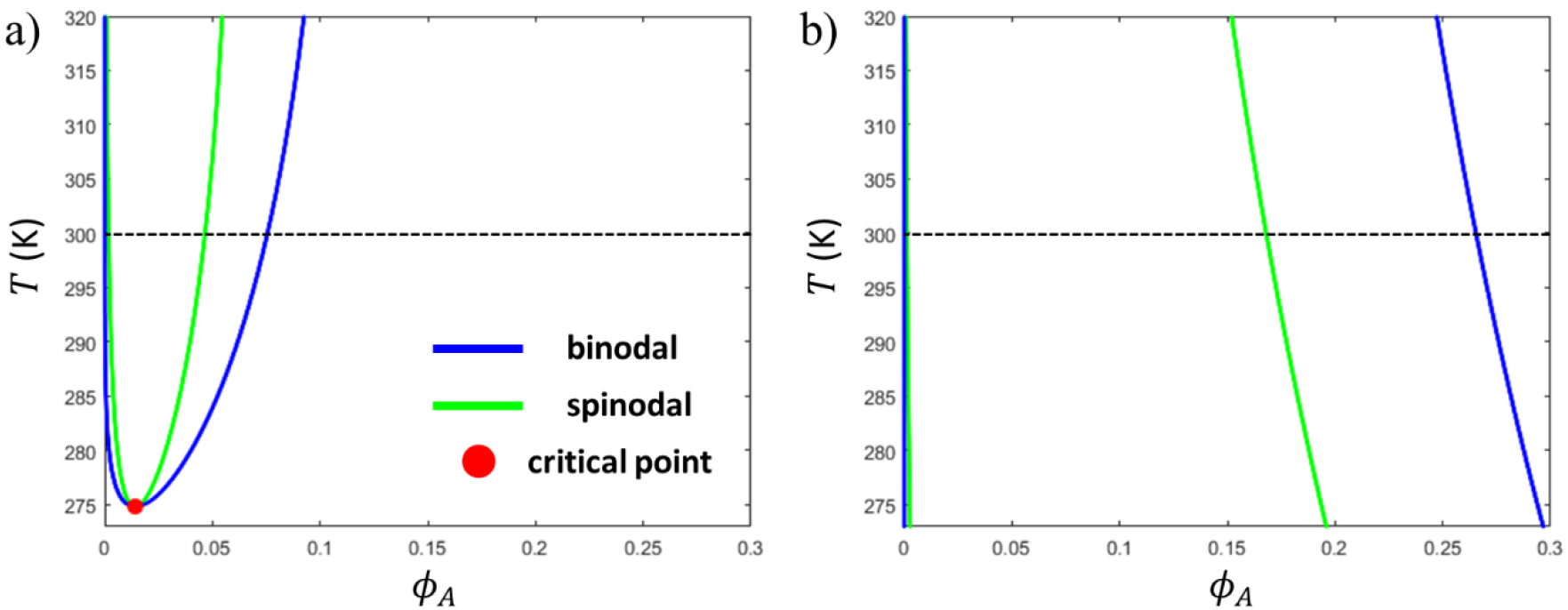
Binary phase diagrams (temperature-composition) calculated for a solution of the scaffold Polymer-A, using input parameters as listed in Table 1, with 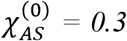 (a) and 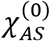 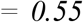 (b). The phase diagram in the (b)-panel represents a small section of an hourglass shaped phase diagram, which lacks a critical point. The horizontal dashed lines indicate *T* = 300 K as the temperature used in most of the ternary calculations.

### Ternary phase behavior: solvent compatibility mediates between associative and segregative LLPS

By means of three sets of calculations of isothermal phase diagrams for the ternary mixture of Polymer-A:Polymer-B:solvent, we demonstrate how non-specific interactions modulate the behavior of Polymer-B between that of an associative client and a species that stimulates the propensity of the scaffold to phase separate, as regularly observed and exploited for crowding agents (44). In these calculations we fix the temperature at *T* = 300 K, except for the second set of calculations, wherein we specifically focus on demonstrating the existence of regions in interaction space for which small temperature changes have a drastic effect on the ternary phase diagram. For technical details concerning the calculation of the ternary phase diagrams, we refer to Section S2 of the SI.

In the first set of ternary calculations, we determine the binodal compositions and the (approximate) position of the critical point(s), as well as the fractions of bound and non-bound stickers as a function of *χ*_*BS*_ and the relative strength of heterotypic sticker association as measured by Δ*ε*. Figures 4a and b plot the obtained isothermal composition diagrams in 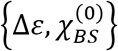-space. The separation between the binodal compositions of the homopolymer solution of the scaffold (see Figure 3a) is reflected by the phase coexistence for *ϕ*_*B*_ = 0. Furthermore, since *χ*_*BS*_ does not exceed its critical value and BB sticker binding is weak, we encounter two-phase coexistence and a single critical point. Since this work is primarily concerned with investigating the size and shape of coexistence regions, as well as quantifying the coexisting compositions, we omit the spinodal curves to optimize clarity in the graphs.

**Figure 4.**
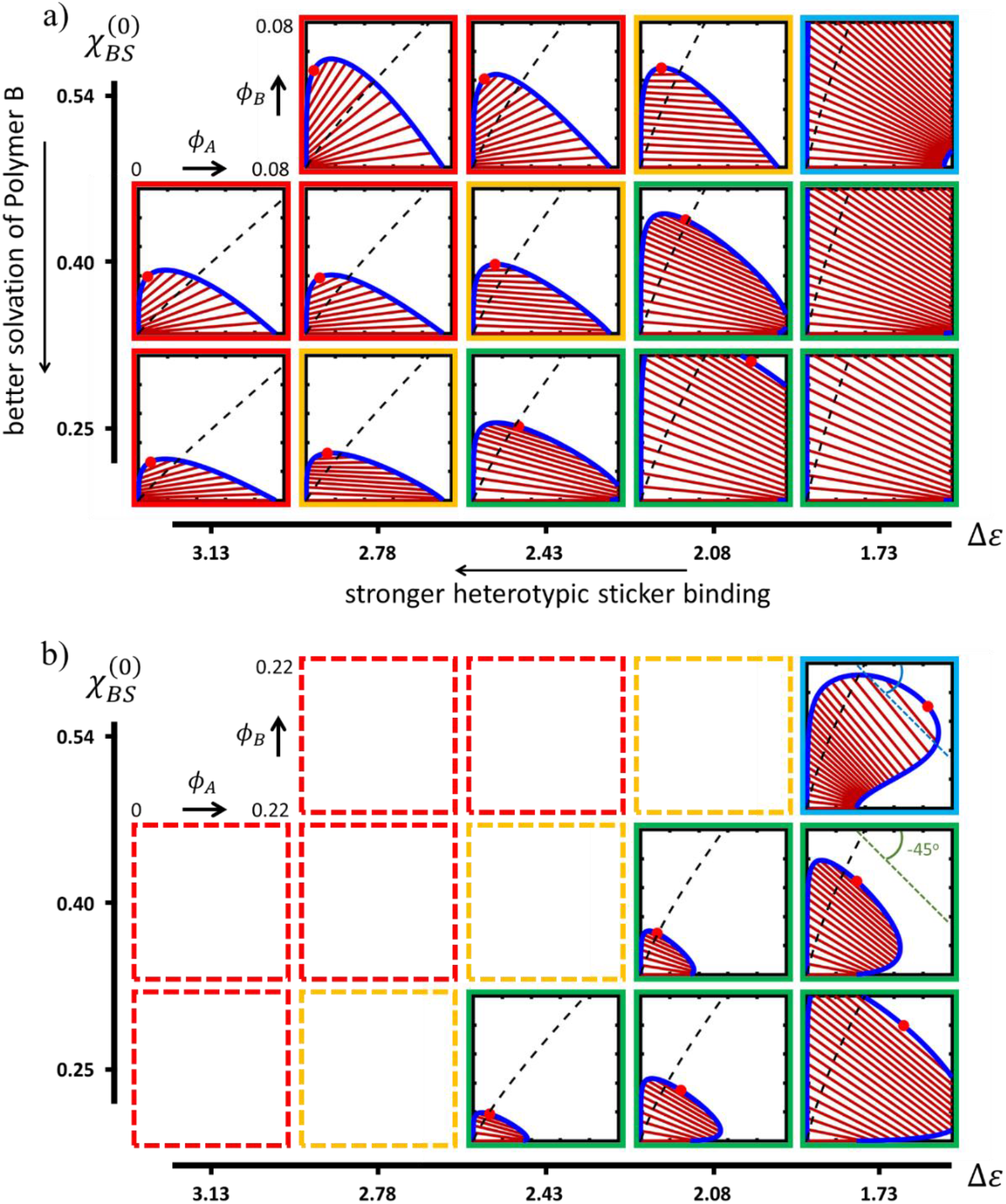
Isothermal (*T* = 300 K) ternary phase diagrams (red symbol: critical point, blue line: binodal, brown lines: tie-lines), plotted as a function of the relative strength of the heterotypic (AB) sticker association and solvent quality for Polymer-B with 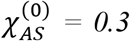. The red, orange, and green/blue frames indicate the regimes of associative, neutral and segregative LLPS, respectively. The dashed black lines indicate the compositions for which 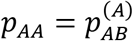. Panel b) shows the phase diagrams of the segregative regime on a more extended composition scale.

The most striking features of the map in Figure 4 are the trends in the size and shape of the miscibility gap, as well as the flip in the (mean) tilt angle of the tie-lines that connect the compositions of the coexisting phases. The tie-line patterns show that for strong heterotypic binding (high Δ*ε*) the two polymers collect in a concentrated phase, giving a positive tilt angle, or, at low Δ*ε*, predominantly assemble in the separate phases, giving rise to a negative tilt. These two modes are dubbed associative and segregative LLPS, and are in Figure 4 indicated by the red and green frames, respectively. In the associative regime, Polymer-B is “recruited” in the concentrated phase of a phase-separated solution of Polymer-A, whereas in the segregative regime Polymer-B amplifies the tendency of Polymer-A to demix, despite the fact that both the specific and non-specific interaction between the polymers are favorable. The transition between the two regimes is marked by a region characterized by a vanishing mean average tilt angle, *i.e.* where Polymer-B partitions approximately equally between the coexisting phases (yellow frames). The division of composition space into regions where homotypic or heterotypic binding is dominant, respectively right and left of the dashed black lines, is discussed below.

A very similar cross-over from associative to segregative LLPS has been observed in Gibbs ensemble simulations of patchy particles that represent protein and RNA components in an implicit solvent (50). Furthermore, more advanced models that include monomer sequence-specificity have shown associative LLPS and segregative LLPS to result, respectively, from similar and dissimilar sticker distribution patterns on both polymers (62, 63). Considering the above, this suggests that the similarity in the monomer sequences of such polymers may well be parametrized by the relative strength of homo- and heterotypic association of stickers on simplified, effective chains that lack sequence specificity. This could, for instance, be of advantage in view of reducing model complexity, computational cost and/or to quantify general thermodynamic parameters (13, 64).

Strongly associative behavior, *e.g.* for 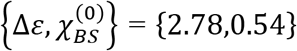 entails a pronounced anticlockwise rotation of the tie-lines at a low but increasing Polymer-B content. This “fanning” expresses a buffering of the composition of the dilute phase. At higher fractions, the concentration of Polymer-B in the dilute phase rises, due to the fact that the regulator species is accommodated relatively well by the solvent and does not exhibit LLPS by itself. Generally, the miscibility gap remains small in the associative regime as strong heterotypic sticker binding stimulates mixing, although, interestingly, an increase is observed with Δ*ε*. In contrast, upon entering the segregative regime (green frames) by decreasing Δ*ε*, *i.e.* moving from left to right in interaction space, the miscibility gap expands rapidly when the homotypic (AA) sticker interaction starts to dominate. To demonstrate this expansion more clearly, we have reproduced the phase diagrams belonging to the segregative regime on a larger composition scale in Figure 4b. As in this regime, the addition of Polymer-B stimulates LLPS in a way similar to what has been observed experimentally for certain crowding agents (44), the two branches of the binodal initially diverge upon increasing the mean concentration of Polymer-B. At elevated fractions of Polymer-B, however, they converge towards a critical point due to fact that the polymers are miscible in the melt. In this case, the tie-lines rotate clockwise, adopting a negative slope with the point of buffering on the concentrated, Polymer-A-rich branch.

Convergence or divergence of the two branches of the binodal in, respectively, the associative and segregative regimes is sometimes referred to as “destabilization” and “stabilization” of the scaffold-rich phase by the addition of the regulating client species (20, 50, 65). Although such terminology conveniently describes the phenomenology, we stress that the phase behavior is a consequence of the free energy density of the complete mixture and not specifically due to the presence of a particular component or interaction between a subset of the components. It is hence illustrative that for the present system the slope in the tie-lines is not solely determined by the relative strength of sticker association, but strongly modulated by the quality of the solvent. Figure 4a shows that if the medium accommodates Polymer-B less well (increasing *χ*_*BS*_), the LLPS becomes associative as the system attempts to reduce polymer-solvent contact.

Furthermore, where in the associative regime the coexistence region increases with *χ*_*BS*_, the opposite trend is observed in the segregative regime. This becomes apparent when comparing for instance the phase diagrams calculated for Δ*ε* = 2.78 with those calculated for Δ*ε* = 2.08. An explanation is that in case of associative LLPS the system lowers its free energy most effectively by i) depleting Polymer-B from the dilute phase and ii) allowing for significant heterotypic association in the concentrated phase. Vice versa, if in the segregative regime the solvent compatibility of Polymer-B becomes better, it is energetically favorable to maximize solvent:Polymer-B contacts, while at the same time enhancing the extent of homotypic A-sticker association via enrichment of Polymer-A in the coexisting phase.

### “Double-reentrant” and dualistic phase behavior in the regime of segregative LLPS

Interestingly, for weak heterotypic sticker association and even higher *χ*_*BS*_ (*i.e.* around *θ*-conditions), a secondary segregative subregime emerges, which we indicate with the blue frame in Figures 4a and b (top right panels). Drastic changes in the trends in size and shape of the miscibility gap, as well as the tie-line slope are observed, when compared to the primary segregative subregime discussed above (green frames). Upon increasing *χ*_*BS*_ at Δ*ε* = 1.73 (rightmost column Figure 4b), the size of the miscibility gap becomes larger again, implying the presence of a minimum. Concomitantly, the divergent behavior of the binodal branches sets in at a higher mean fraction of Polymer-B. The Polymer-A rich branch initially curves upwards and away from the horizontal axis. The tie-lines rotate in a clockwise manner around the buffering point on the concentrated branch, however now reaching a maximum slope that is steeper than −45°, exceeding the maximum slope observed in the primary segregative subregime. Hence, in this secondary segregative subregime a region seems to emerge in composition space where the Polymer-B-rich becomes more concentrated than a coexisting Polymer-A-rich phase. The deviant shape of the miscibility gap, in combination with the marked change in the tie-line slope no longer seem to express a singular behavior, but rather indicates the presence of two coexistence regions, that for the given input seem to overlap. Since two coexistence regions in an asymmetric ternary phase diagram likely exhibit a different temperature dependence, we perform a second set of calculations (see Figure 5), where we i) implement minor temperature variations and ii) extend the range for *χ*_*BS*_ to demonstrate the presence of multiple coexistence regions and to probe the extent of this secondary segregative subregime.

**Figure 5.**
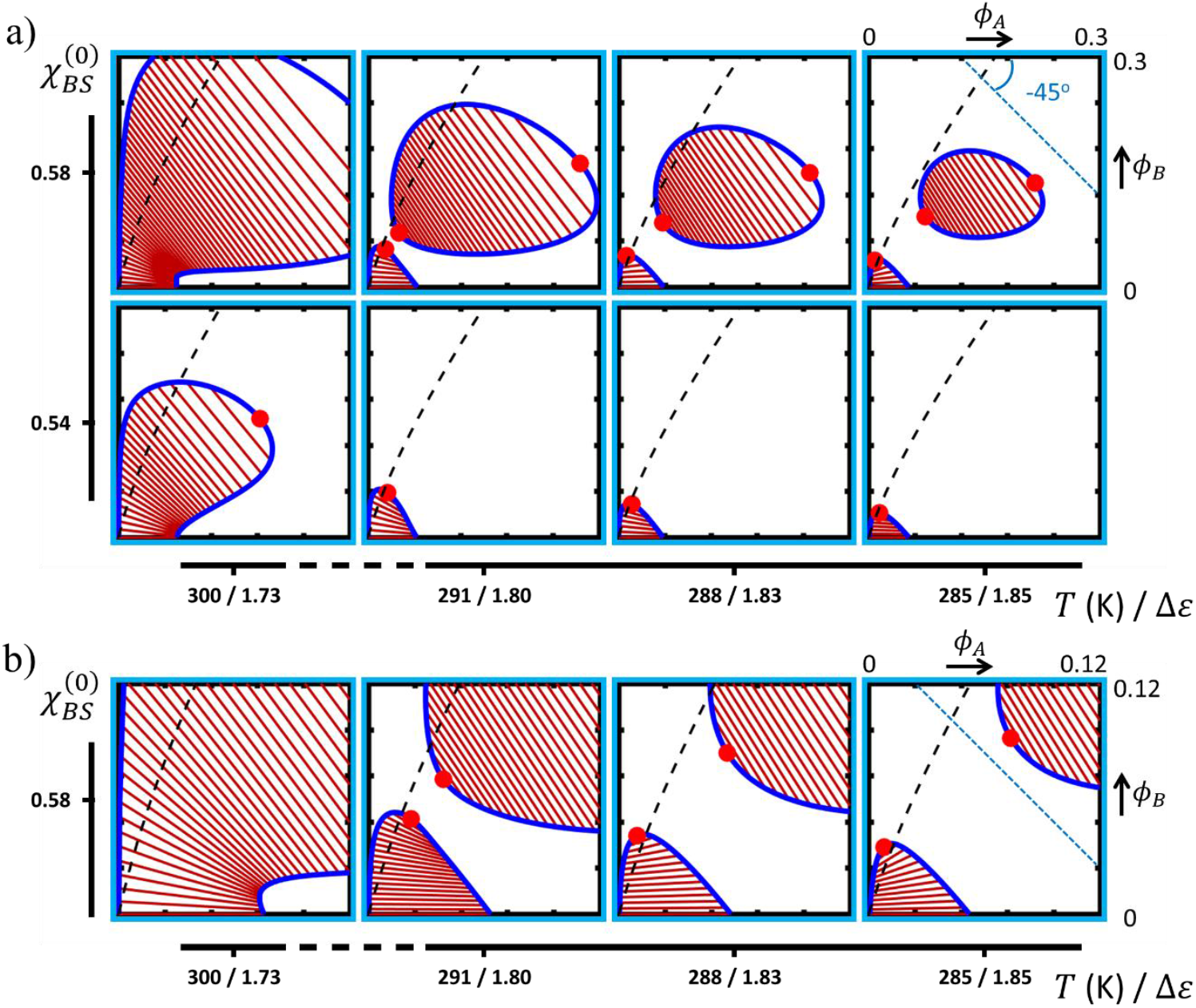
Isothermal ternary phase diagrams (red symbols: critical points, blue lines: binodal, brown lines: tie-lines), plotted as a function of the absolute temperature and 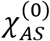 in the secondary subregime of segregative LLPS, with 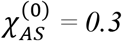. The relative strength of heterotypic sticker association (Δ*ε*) has been indicated for each temperature. The dashed black lines indicate the compositions for which 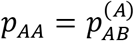. Panel b) is a magnification of the top row of the a)-panel to facilitate visualization of the tie-line patterns.

Figure 5a (top row) indeed shows that if *χ*_*BS*_ further increases, a slight decrease in the temperature results in a clear separation into two miscibility gaps, giving rise to three critical points: one associated with the primary coexistence region due to the LLPS of the scaffold solution and two with a second, elliptical region appearing at elevated polymer concentration. Since such a closed loop binodal by itself represents reentrant phase behavior (18, 37, 38, 58, 66, 67), the model in fact predicts “double-reentrant” behavior with the simultaneous presence of the primary coexistence region. It would be an interesting challenge to validate this behavior for solutions of biomacromolecules. The bottom row of Figure 5a shows that for a somewhat lower *χ*_*BS*_ (*i.e.* Polymer-B being slightly better accommodated by the solvent), the looped miscibility gap contracts rapidly with decreasing temperature, leaving only the primary coexistence region.

The size of the two miscibility gaps both decrease with temperature, despite the weakening of heterotypic sticker binding and increase in the solvent-polymer interaction parameters. This expresses the dominance of homotypic AA sticker association in determining the phase diagram. Interestingly, the tie-lines associated with the closed loop binodal all have a slope steeper than −45° and fanning in the pattern is near absent. The latter is in conjunction with the fact that the tangent lines at the two critical points are nearly parallel. Hence, for the looped binodal, independent of the overall composition, the Polymer-B-rich phase is more concentrated than the Polymer-A-rich phase, which suggests that the prioritization implied by terms such as “client”, “crowder” and “scaffold” may lose meaning depending on the relative contributions of competitive sticker binding and solvent compatibility. In contrast to the closed loop coexistence region, the tie-line slope associated with the primary miscibility gap remains less steep than −45° and even becomes positive at *T* = 285 K (see Figure 5b). In other words, as the temperature decreases, we observe a crossover from segregative to associative LLPS, as in the present case AA association has a stronger temperature dependence than heterotypic binding. The effect becomes clear if we calculate the sticker exchange energy Δ*ε* (indicated on the horizontal axis in Figures 5a and b), which shows an inverse relation with temperature. Hence, segregative and associative LLPS can in principle be encountered within the same phase diagram, here associated with two miscibility gaps.

### Homotypic versus heterotypic sticker association

Besides the binodal compositions and critical points, our calculations also produce the fractions of bound (*p*_*ij*_) and non-bound (*p*_*i*_) stickers of each of the two polymers in the coexisting phases. Quantification and prediction of these fractions is of high interest, as not only the number of bound stickers per chain, but also the partitioning between homotypic and heterotypic complexes directly affects the constitutive properties (viscosity/viscoelasticity) of a droplet phase, as well as the diffusive dynamics of its components. For the interaction matrix used in the present study a concentration-dependent competition between homotypic and heterotypic binding applies specifically to the scaffold Polymer-A. For the client Polymer-B, due to the very weak BB association it is imperative that bound B-stickers are almost exclusively involved in heterotypic association, irrespective of composition.

To show how the fractions of non-bound and bound A-stickers depend on the binodal compositions, we have reproduced an illustrative selection of the phase diagrams in Figures 4 and 5 in a 3D plot (Figure 6), of which the (x,y)-plane represents composition space and the z-axis quantifies the fractions of bound and non-bound A-stickers. In all ternary phase diagrams (Figures 4-6) we have divided (dashed black lines) composition space in the regions where either AA-homotypic or AB-heterotypic sticker association dominates. Which complex dominates becomes clear in Figure 6, which shows the coincidence of the crossover of the green (*p*_*AA*_) and red 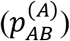 curves with the point where the dividing line intersects with the binodal. The dividing lines have been obtained analytically by solving the sticker binding model, subject to the constraint: [*AB*] ≡ *K*_*AB*_[*A*][*B*] = 2[*A*_2_] = 2*K*_*AA*_[*A*]^2^, here formulated on the basis of molar concentrations to illustrate the equivalence of the free energy model with a “chemical” description of the binding equilibria. This constraint allows for expressing the concentration of non-bound B-stickers [*B*] as a direct function of [*A*]. Substitution in the mass balance Equations:

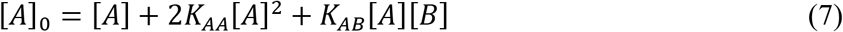

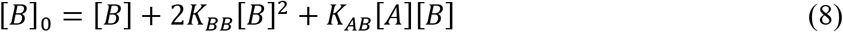

gives the relation between the mean volume fractions for which 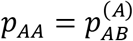:

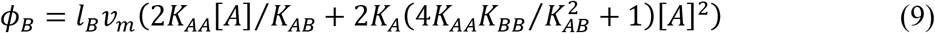

with: 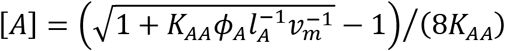, and *v*_*m*_ = 1 mol l^−1^ a reference molar volume.

**Figure 6.**
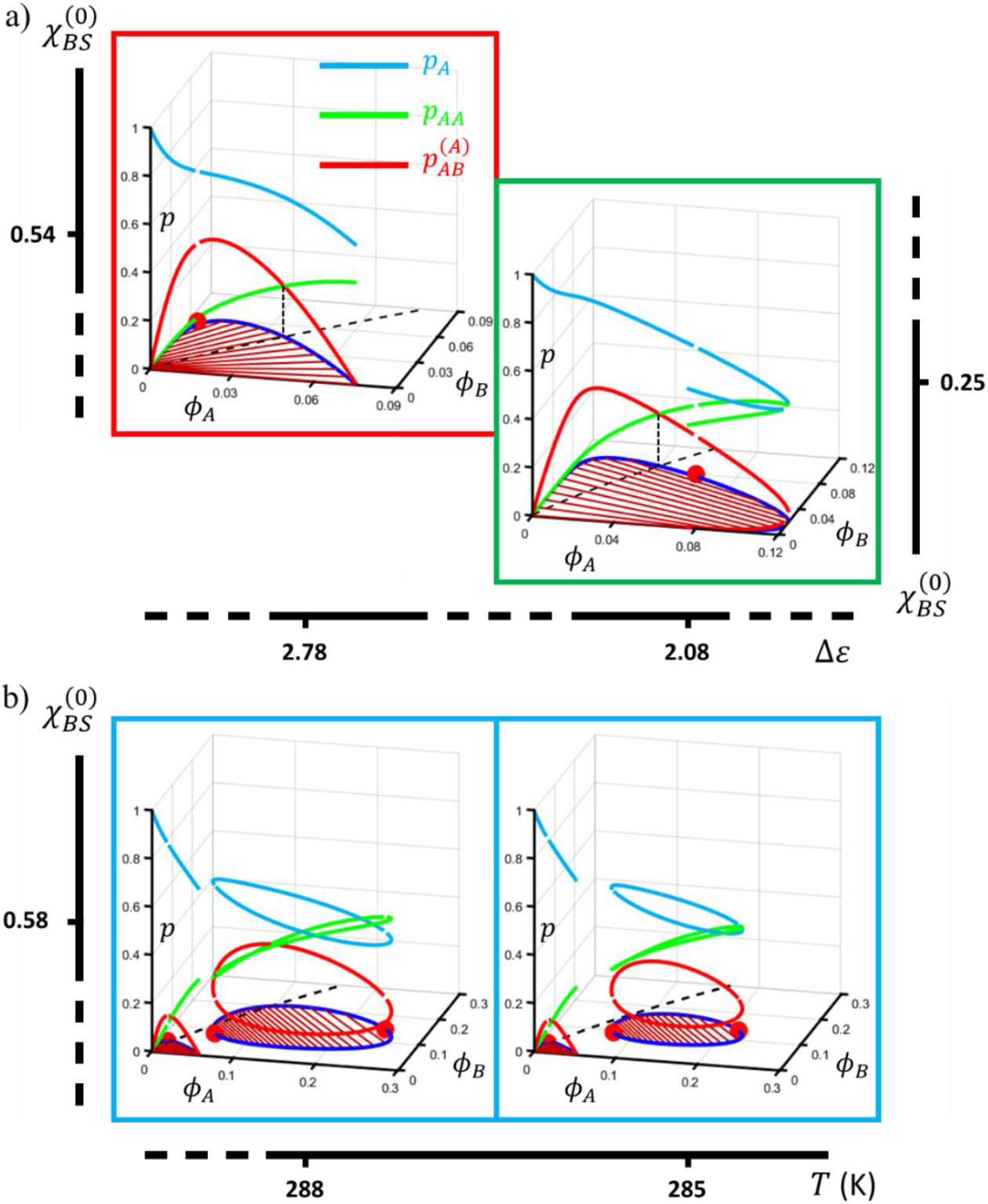
Exemplary 3D representations of the relation between the phase diagram (binodal and critical points) and the fractions of bound and non-bound A-stickers for a) the associative (red frame) and primary segregative subregime (green frame), as well as b) the secondary segregative subregime (blue frames). The ternary phase diagrams in a) and b) have been reproduced from Figures 4 and 5, respectively. The dashed black lines represent Equation (9) and indicate the compositions for which 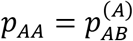.

Figures 6a and 4 show that the average slope of the dashed dividing line in the composition-plane exhibits a reciprocal relation with Δ*ε*: as heterotypic association becomes stronger, the line rotates clockwise around the origin. The fact that the critical point and the tie-lines rotate in the opposite direction leads to a rich variation in the association pattern of the A-stickers. If the dividing line crosses the coexistence curve exactly at the critical point, which for the present input is the case for weakly segregative LLPS, homotypic A-sticker binding dominates at the Polymer-A-rich branch of the binodal and heterotypic binding is more prominent at the Polymer-B-rich branch. Hence, in this scenario the preference for a particular type of association in either of the coexisting phases does not depend on the overall composition. In most cases however, the dividing line intersects one of the binodal branches, as shown in Figure 6, which means that for one of the coexisting phases the overall composition dictates whether homotypic or heterotypic binding dominates. In case of associative LLPS this concerns the concentrated binodal branch, whereas for segregative LLPS (primary subregime) this applies to the more dilute Polymer-B-rich phase.

The way the bound A-stickers partition between homotypic and heterotypic complexes in the secondary segregative subregime (Figure 6b) is similar as far as the primary miscibility gap concerns. Figure 5 shows that upon a slight lowering of the temperature the dividing line rotates clockwise, in favor of heterotypic binding. As for the closed loop miscibility gap, depending on the temperature intersection of the binodal by the dividing line may occur once, twice or not at all. For 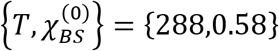 for instance, the concentrated branch is dissected twice (see left panel in Figure 6b), which leads to two regions on this branch where homotypic binding is more prominent than heterotypic binding, despite the fact that this concerns the Polymer-B-rich phase. By contrast, for 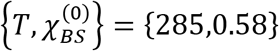, the loop is situated entirely in the region where homotypic A-sticker binding dominates (Figure 6b, right panel).

### Dualistic phase behavior in the regime of associative LLPS

Above we have seen that the interplay between sticker binding and polymer-solvent interaction may lead to a scenario where associative and segregative LLPS are encountered in the same phase diagram (rightmost panel in Figure 5b). In this case, each mode of demixing is associated with a separate miscibility gap. Since our starting point for demonstrating this dualistic behavior was the primary *segregative* regime, the associative behavior, as a matter of speaking, made a “reappearance”. In a third and last set of calculations, we demonstrate that the opposite scenario, *i.e.* a reappearance of segregative LLPS in the associative regime, is consistent with dualistic behavior to occur even within a single coexistence region. We recall that for strong heterotypic binding (high Δ*ε*), the associative nature of the LLPS is particularly strong of if the scaffold Polymer-A is energetically better accommodated by the solvent than Polymer-B (see leftmost column in Figure 4), so *χ*_*AS*_ < *χ*_*BS*_. The explanation for this is that expelling both polymers towards a concentrated phase minimizes interaction between Polymer-B and solvent and maximizes the contribution from heterotypic sticker binding, being the preferred mode of sticker association.

In what follows, we address the inverse situation, *i.e.* wherein Polymer-B is better accommodated by the solvent than Polymer-A, but maintain a high Δ*ε*. We hereto set Δ*ε* = 3.13 and increase 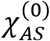 from 0.3 to 0.55, *i.e.* a subcritical value somewhat exceeding *θ*-conditions that, in combination with homotypic A-sticker binding, gives rise to coexistence across the complete temperature range (see Figure 3). We calculate the isothermal ternary phase diagram while varying 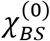 in the range of 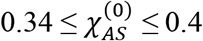 at *T* = 300 K. Figure 7 displays the results. To enhance clarity in showing the changes in the tie-line pattern, we have enlarged the scale of the vertical axis. Figure 7a shows that the tie-line pattern undergoes drastic changes when lowering 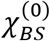. Comparing the top panel of Figure 7a with its counterpart in Figure 4 (leftmost middle panel) reveals that the associative nature of the LLPS of the binary solution initially weakens. The fanning in the tie-line pattern becomes less pronounced and the buffering of the dilute phase, characteristic for associative LLPS, occurs for a higher overall Polymer-B fraction.

**Figure 7.**
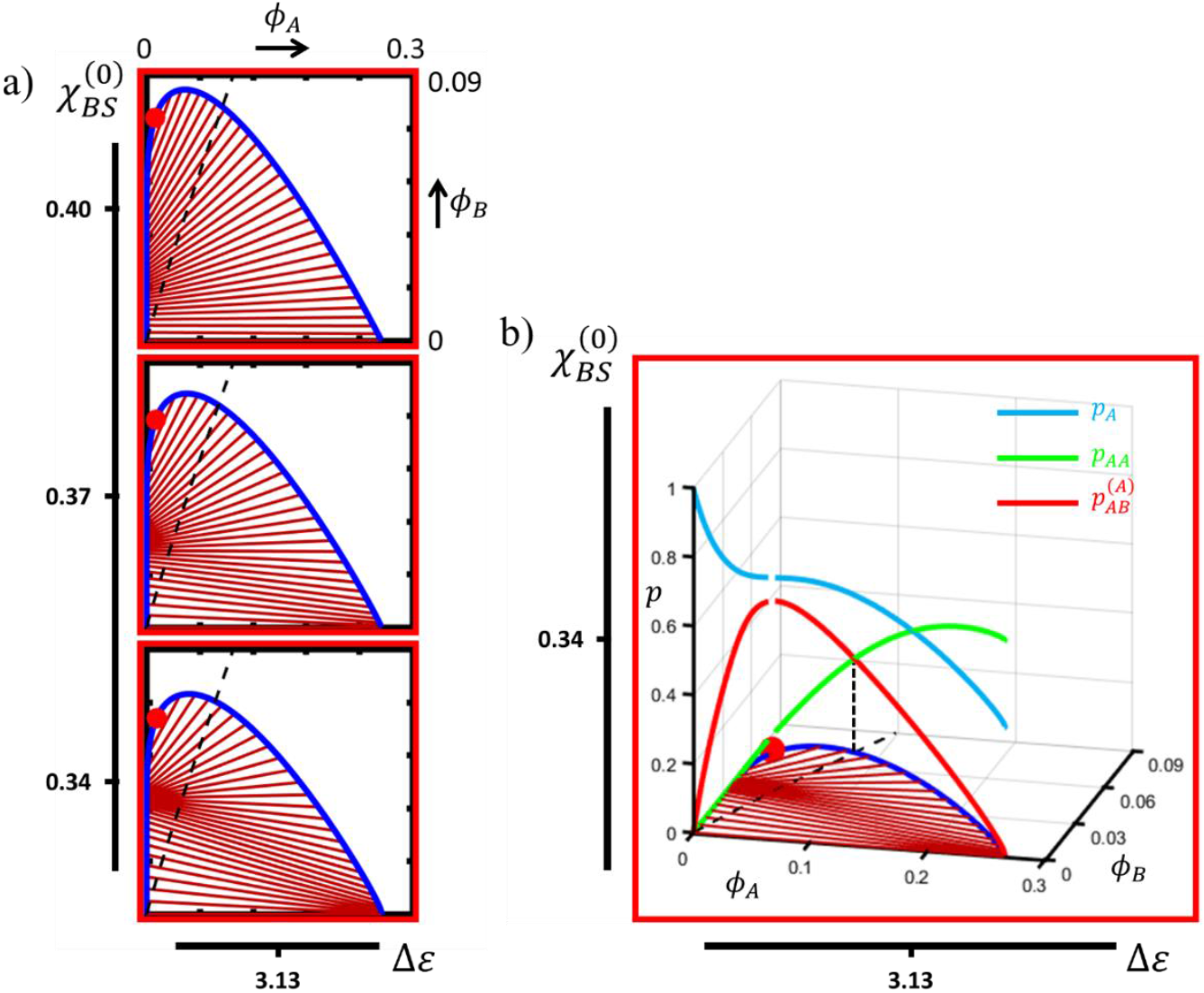
Isothermal (*T* = 300 K) ternary phase diagrams (red symbol: critical point, blue line: binodal, brown lines: tie-lines), calculated for Δ*ε* = 3.13 and 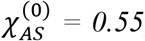, plotted as a function of the solvent quality for Polymer-B (a). For clarity, we have stretched the y-scale. Panel (b) is a 3D reproduction of the bottom panel in a) to visualize the bound and non-bound fractions of A-stickers in composition space. The dashed black lines represent the compositions for which 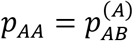.

Reducing 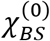 (middle and bottom panel in Figure 7a) even causes a negative tie-line slope at low Polymer-B content, hence expressing features of segregative LLPS despite the high Δ*ε*. Apparently, since at a low fraction of Polymer-B the number of heterotypic sticker complexes and associated energetic gain is low, the most effective way for the system to minimize its free energy is by separating Polymers-A and -B in a, respectively, concentrated and a dilute phase. This way, LLPS maximizes homotypic A-sticker association, as well as contact between the solvent and Polymer-B, while at the same time minimizing Polymer-A-solvent contact. Indeed, the contribution from homotypic A-sticker binding in the concentrated phase is more significant now due to the widening of the miscibility gap of the temperature-composition diagram of the homopolymer solution (Figures 3). Indeed, comparison of Figures 7b and 6a shows that as a result of this, the fraction of A-stickers involved in homotypic binding at low *ϕ*_*B*_ is considerably higher for *χ*_*AS*_ = 0.55 than for *χ*_*AS*_ = 0.3.

As the overall fraction of Polymer-B increases for low *χ*_*BS*_, the contribution of heterotypic sticker association to the free energy becomes more prominent and eventually starts dominating the phase behavior. At this point, the nature of the LLPS changes from segregative to associative, as expressed by the tie-line slope changing from negative to positive. Hence, in the present case both types of LLPS occur within the same miscibility gap, which, for sufficiently low 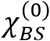 gives rise to two points of buffering, *i.e.* where the tie-lines converge (Figure 7a, bottom panel). In the segregative part of the miscibility gap this concerns the concentrated, Polymer-A-rich phase, whereas in the associative region the dilute phase is being buffered, though at an elevated Polymer-B fraction.

## Conclusions and outlook

Using a ternary mean field stickers-and-spacers model, we demonstrate that the liquid-state phase behavior of a solution of a weakly but multivalently associating scaffold polymer and a client can be very rich due to interference between contributions associated with sticker binding and solvent compatibility. The regulating properties of the client strongly depend on the difference in solvent compatibility of the two polymers. In effect, instead of acting as an inert medium the solvent modulates the properties of the client between that of an associating ligand and a species that stimulates separation of a scaffold-rich phase, as regularly observed for crowding agents. We demonstrate how the interplay between competitive sticker association and polymer-solvent interactions subdivides parameter space into multiple regimes for associative and segregative ternary LLPS. Interestingly, for weak heterotypic sticker association, the phase diagram exhibits two miscibility gaps if the client species is solvated relatively poorly. Depending on the temperature, the two coexistence regions may combine segregative and associative behavior within the same ternary phase diagram. Vice versa, if the solvent accommodates the client better than the scaffold, such dualistic phase behavior can even occur within a single miscibility gap.

This modeling study shows that although specific binding is generally responsible for the phase behavior of solutions of associating biopolymers, the solvent, even if providing for a reasonably well accommodating environment, mediates the role specific components play in determining the phase diagram. The *in-vivo* implication of this is that mutations in chain segments that do not take part in specific binding may have a pronounced effect on bioregulatory processes through affecting MLO composition and stability. Of course, although all input is physically and physiologically reasonable, the results of this purely computational study require validation. This particularly concerns the predicted dualistic and “double reentrant” phase behavior. With this respect, we note that it is not imperative that a homopolymer solution of the scaffold exhibits an LCST, as in this work. The calculations show that the isothermal ternary behavior is a consequence of the relative strength of various types of interactions, which may well follow similar trends in case of a UCST.

## Acknowledgements

The authors acknowledge financial support by the German Research Foundation (Priority Program 2191, Molecular mechanisms of functional phase separation, Project 419138152) and no conflict of interest.

## Author contributions

J. J. M. performed the research and wrote the manuscript. All other authors assisted in defining the model and contributed to the writing. E. A. L., S. H. P. and J. J. M. secured the funding.

## SUPPLEMENTARY INFORMATION

### S1 Model derivation

The dimensionless Helmholtz mixing free energy per mer or lattice site for a solution of two associating polymers A and B in a non-associating solvent S is given by:

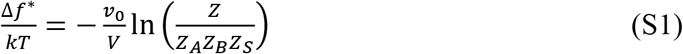

Here, *v*_0_ is the fundamental monomeric site volume and *V* the total volume of the incompressible mixture. *Z* and *Z*_*k*=*A,B,S*_ are the partition functions of the mixture and pure species. We write the partition function for the mixture in absence of non-specific exchange interactions as (S1, S2):

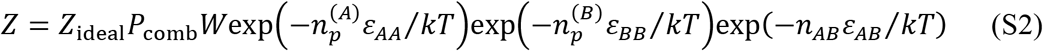

Here, *Z*_ideal_ is the ideal gas partition function, *ε*_*ij*_ the binding energies of homotypic and heterotypic complexes, 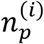 the number of homotypic complexes (paired stickers) of type *i* and *n*_*AB*_ the number of heterotypic complexes. *P*_comb_ counts the number of ways 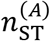 and 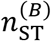 stickers can be combined into 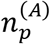 and 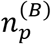 homotypic and *n*_*AB*_ heterotypic complexes:

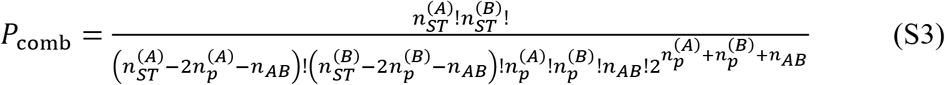

and *W* the probability that all stickers involved in binding can be found close enough to their partners to form bonds in the absence of attractive forces:

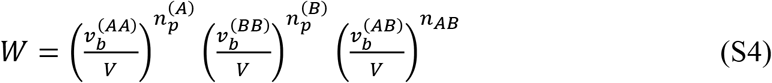

with 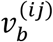 the bond volumes, which for simplicity we all set to: 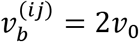.

The resulting ideal and sticker contributions to the normalized free energy density of the mixture are:

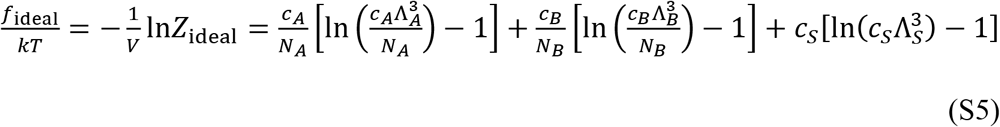

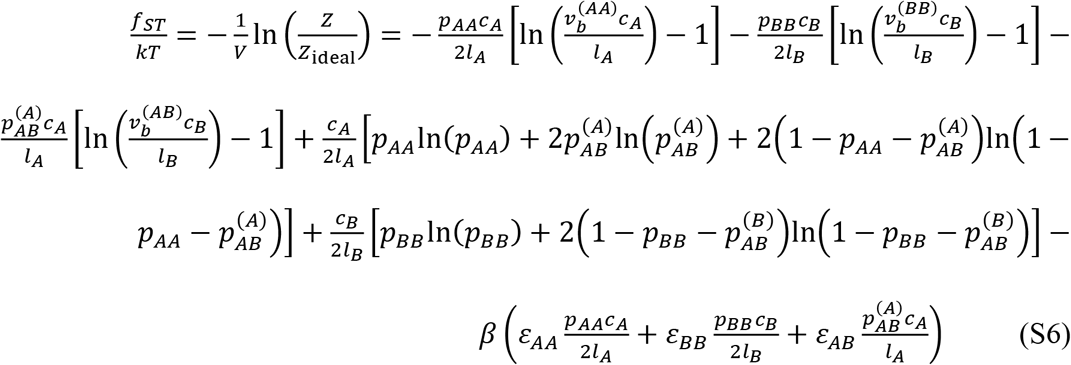

where the molecular sizes have been normalized by that of a solvent molecule giving *N*_*i*=*A,B*_ as effective degree of polymerization in mers with volume *v*_0_. *c*_*k*_ is the number density of *k*-monomers and Λ_*k*_ is the thermal wavelength. In Equation (S6), *p*_*ii*_ is the fraction of bound *i*-stickers involved in homotypic *ii*-complexes. 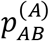 and 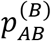 are the fraction of *A*-stickers and *B*-stickers involved in heterotypic complexes, respectively. The latter are related via: 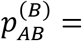 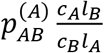, with *l*_*i*_ the (average) number of monomers between *i*-stickers. Applying the equilibrium condition for sticker binding 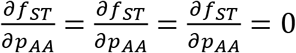 yields:

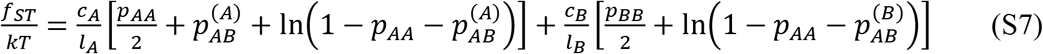

as well as Equations (4)–(6) in the main text. Equivalently, we obtain for the free energy density of the pure states:

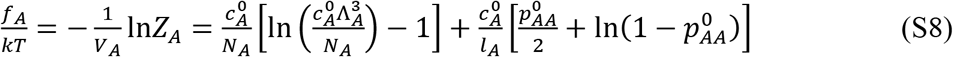

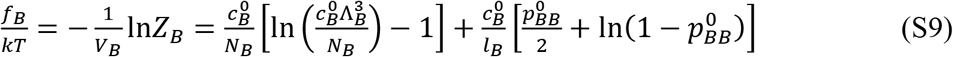

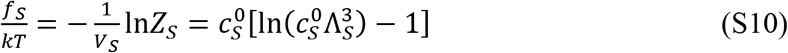

with 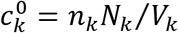 the monomer density in the pure state. Equations (S1), (S7)–(S10) give together with the incompressibility constraint *V* = *V*_*A*_ + *V*_*B*_ + *V*_*s*_ = *v*_0_(*n*_*A*_*N*_*A*_ + *n*_*A*_*N*_*A*_ + *n*_*s*_):

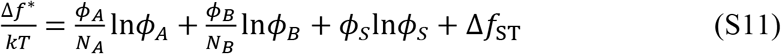

of which the last term is given by Equation (3) in the main text. We obtain the total dimensionless free energy per monomer Δ*f* (Equation (1)) by adding the mean-field non-saturating exchange interactions arising from mixing and parametrized by the Flory interaction parameters.

### S2. Procedure for calculating ternary phase diagrams

We assume a ternary mixture consisting of two sticky polymers A and B in a solvent S. To calculate the ternary phase diagram we add small increments of Polymer-B to a mixture of Polymer-A and solvent, or remove small quantities of Polymer-B from the ternary mixture, and calculate the resulting binodal compositions. To determine the composition of two coexisting binodal phases *α* and *β* requires numerical calculation of the four volume fractions 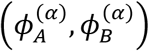 and 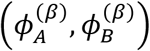, the volume fractions of solvent being dependent and obtained via the incompressibility assumption: 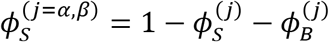. In principle, equalizing the exchange chemical potentials and osmotic pressures of components A and B in the coexisting phases gives these values, for instance via a tetravariate Newton-Raphson (NR) procedure. However, the multivariate nature of the problem, in combination with the complexity of the free energy with its contributions from specific and non-specific interactions compromises the ability of a single root finding procedure to converge. Instead, we take a more robust approach based on a reduction of the number of variables and splitting the problem into a set of nested bi- and monovariate NR loops.

To find the binodal compositions for a given overall composition 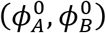 in the miscibility gap (see Figure S1) we follow the following iterative procedure. We i) map the composition coordinates onto a single coordinate *h*(*ϕ*_*A*_, *ϕ*_*B*_) running along a cross-section at a given angle *θ* as indicated in Figure S1, ii) find *h*^(*j*=*α,β*)^ and hence the compositions 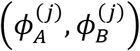 at which this line intersects both branches of the binodal, and iii) optimize the value for *θ* so as to fulfill the binodal criterium. We use a bivariate NR loop to find estimates for *h*^(*j*)^, embedded within a main loop defined by a monovariate NR procedure that converges *θ* to the value that represents the actual tilt angle of the tie-line. Once that value has been reached, *h*^(*α*)^ and *h*^(*β*)^ represent the compositions of the coexisting phases and not merely two unassociated binodal concentrations.

**Figure S1.**
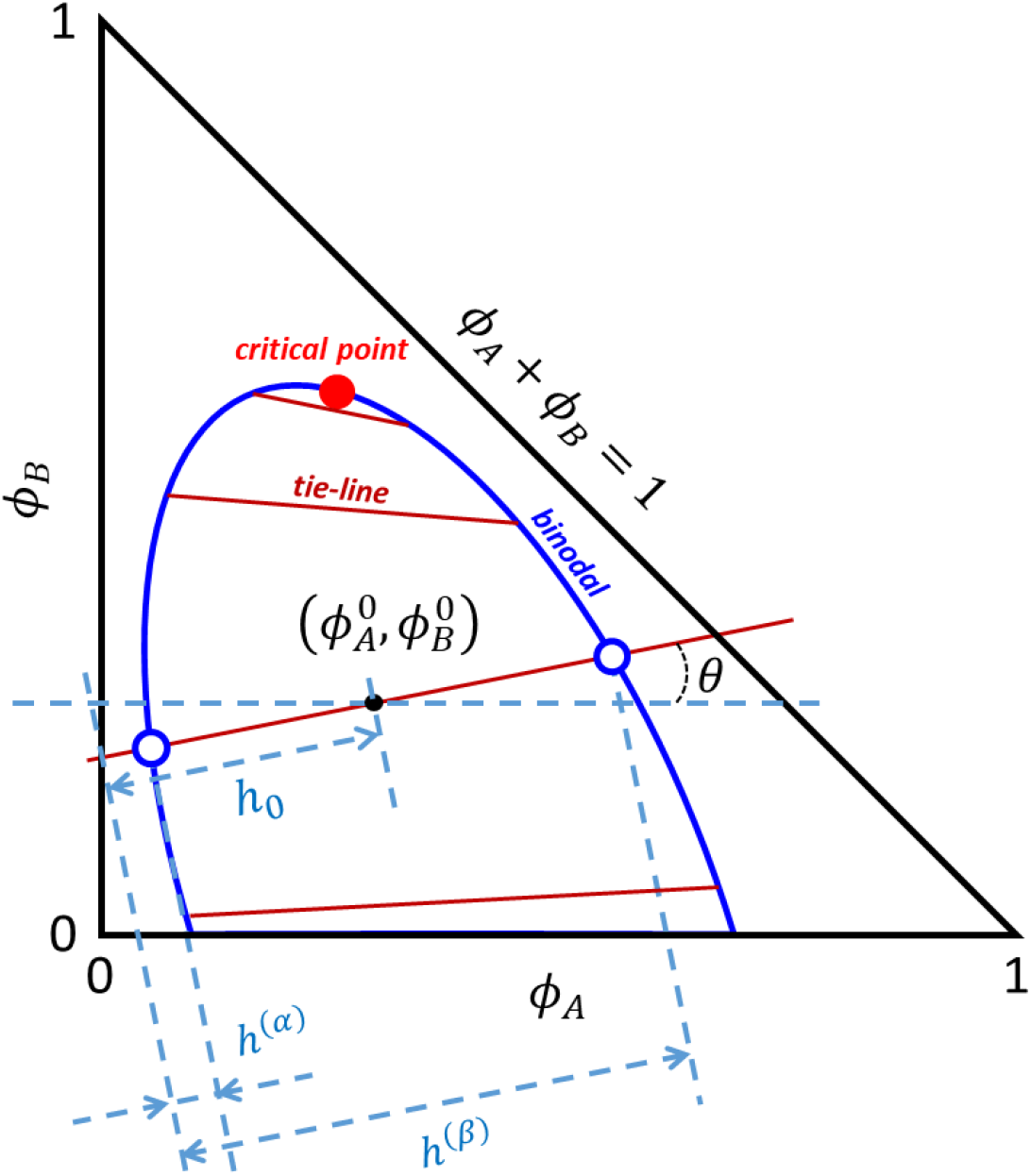
Schematic ternary phase diagram drawn within the plane determined by the independent volume fractions *ϕ*_*A*_ and *ϕ*_*B*_. The composition of the coexisting *α*- and *β*-phases, indicated by the open symbols, are obtained by finding the value for the tie-line tilt angle *θ* that minimizes the difference in the chemical potential and the osmotic pressure between the two phases.

Before providing the equations associated with the NR loops, we give the relation between *h* and *ϕ*, as defined by straightforward goniometric arguments (see Figure S1):

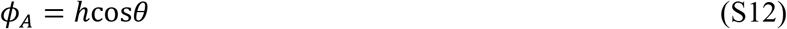

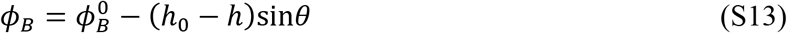

Depending on the value for *θ*, the upper bound for *h* is given by the following condition:

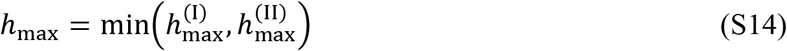

with

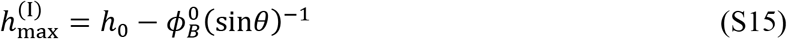

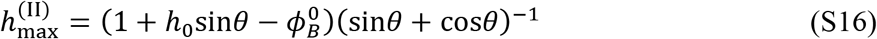

In all calculations in this work, |*θ*| is sufficiently small, so that the lower bound for *h* is always *h*_min_ = 0. To find *h*^(*α*)^ and *h*^(*β*)^, we numerically approximate the roots of the following system of equations using the (inner) bivariate NR procedure:

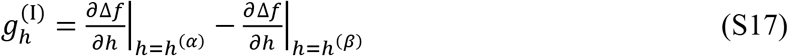

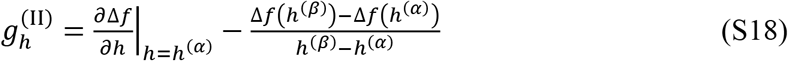

These conditions are equivalent to those for finding the coexisting compositions in a binary mixture, where *h* = *ϕ*_*A*_ ≡ *ϕ* and *θ* is zero per definition. In effect, Equations (S17) and (S18) assure fulfillment of the prerequisite of equal osmotic pressure. The iteration of the NR routine proceeds according to:

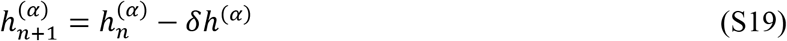

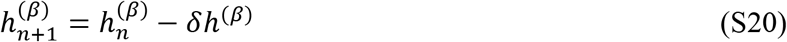

with *n* indicating the current iteration and

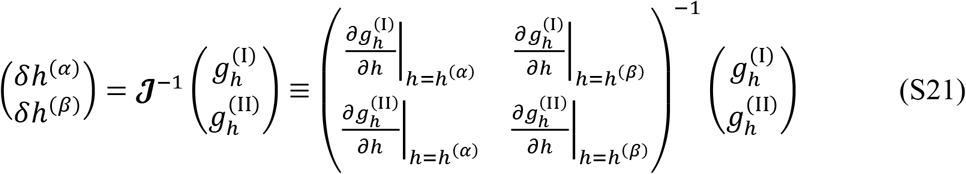

Evaluation of the elements of the Jacobian matrix 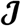 requires the first and second derivative of the free energy with respect to *h*, which, using the chain rule and Equations (S1) and (S2), we write as a function of the free energy derivatives with respect to the independent volume fractions:

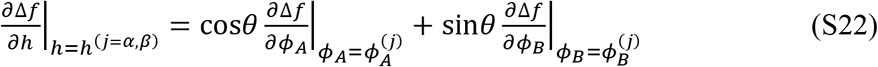

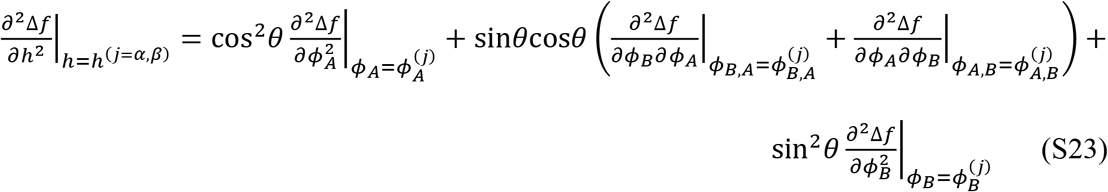

The tilt angle *θ* of the actual tie-line through the point 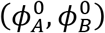 is a value for which the following condition is met, owing to the exchange chemical potentials being the same in the coexisting phases:

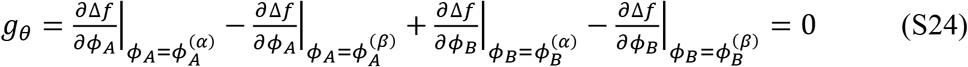

The decrement *δθ* of the monovariate (outer) NR procedure that optimizes the value for *θ* is given by: 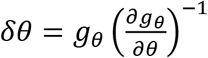. In order to evaluate the derivative 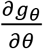 we require:

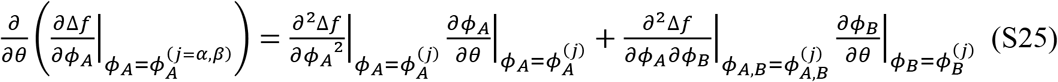

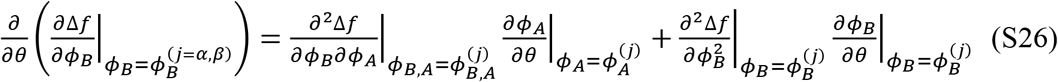

We have:

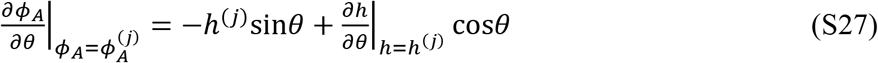

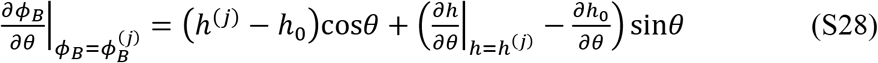

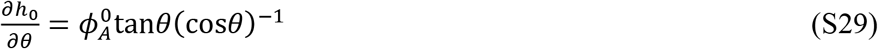

The derivatives 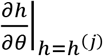 are obtained numerically using the central difference approximation:

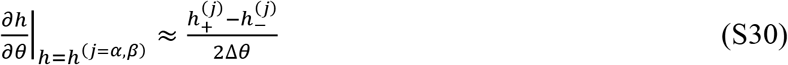

with Δ*θ* a small variation in the current value for *θ*, and 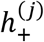 and 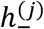 the values for *h* corresponding *θ* + Δ*θ* and *θ* − Δ*θ*, obtained using the bivariate NR procedure outlined above.

To allow for maximal flexibility and numerical stability, we evaluate the convergence criteria for the various NR routines as independent input parameters. The same holds for the increment in 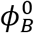 and Δ*θ*, of which the former determines the total number of calculated binodal compositions. For each subsequent point, we evaluate 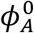 to be the average of the volume fractions in the coexisting phases calculated in the previous step. For simplicity, we do not explicitly calculate the critical point but approach its position very closely, taking such small increments in the overall composition that the two branches of the binodal almost connect.

